# Intestinal tissue response to Shiga toxin exposure

**DOI:** 10.1101/2024.06.11.598485

**Authors:** Kendal L. Hanson, Alison Ann Weiss

## Abstract

Shiga toxin (Stx) is produced by some pathogenic strains of *E. coli*. To study the impact of Stx on the human intestine we utilized human intestinal organoids (HIOs) and human intestinal enteroids (HIEs) grown as human intestinal enteroid monolayers (HIEMs) in transwells. To establish optimal experimental conditions, HIEMs were grown with or without mesenchymal cells added to the basolateral wells to recapitulate the interactions between the intestinal epithelium and underlying mesenchyme. Monolayer barrier integrity was determined through transepithelial electrical resistance (TEER) readings. Apical saline was used on the apical surface since growth medium caused uneven development of the TEER. Medium used for epithelial cells contains added growth factors, while mesenchymal medium lacks these growth factors. We have shown that mesenchymal cells can maintain the epithelial monolayer in medium lacking growth factors suggesting they produce these factors. Furthermore, growth factors produced by mesenchymal cells need to buildup in the medium over time, since daily medium changes were not as effective as medium changes performed every three days. We have also shown that addition of growth factors is toxic to mesenchymal cells. Epithelial cells were more resistant to Stx2 than the mesenchymal cells, and mesenchymal cells contributed to epithelial cell death. Epithelial cells tolerated luminal exposure better than basolateral exposure. These studies demonstrate the importance of understanding tissue interactions in a disease state when using in vitro and in vitro models.

**Importance:** These studies have cemented the need for complex cell culture models when studying host-pathogen interactions. Common animal models such as mice are resistant to *E. coli* O157:H7 infections and intestinal delivery of Stx2, while humans appear to be sensitive to both. It has been proposed that in humans, STEC-mediated intestinal damage destroys the intestinal barrier and allows basolateral access to Stx2. In mice, there is no epithelial damage, therefore they are resistant to epithelial delivery of Stx2, while remaining sensitive to Stx2 injection. Our studies show that like mice, the human epithelial layer is quite resistant to Stx2, and it is the sensitivity of the mesenchymal cells that kills the epithelial cells. We have shown that Stx2 is transported through the intact epithelium without causing damage to the resistant epithelial layer. Understanding tissue interactions during infections is therefore critical in determining the effects of pathogens on human tissues.

## Introduction

*E. coli* O157:H7 is a pathogenic strain of *Escherichia coli* that is characterized by Shiga toxin (Stx) production. Due to its Stx production and association with bloody diarrhea, it is often known as Shiga toxin producing *E. coli* (STEC) or enterohemorrhagic *E. coli* (EHEC). Found in the bovine intestinal tract, it is a common foodborne pathogen spread by contaminated produce, undercooked meat, and dairy products. It is known to cause bloody diarrhea and hemolytic uremic syndrome in humans [1], and according to the CDC, is responsible for 40% of the approximate 265,000 illnesses and 100 deaths from *E. coli* infections each year in the US alone [2]. Of those diagnosed with STEC infections, 5-10% develop hemolytic uremic syndrome (HUS) [3]. This disease primarily affects children and is characterized by microangiopathic hemolytic anemia, thrombocytopenia, and potentially renal failure and death [4].

Stx is a bacterial toxin with an AB_5_ structure. The A subunit is the active portion, and the five B subunits make up the binding portion of the toxin. The A-subunit is enzymatically cleaved into the larger A1 subunit which possesses the toxic activity, and a smaller A2 subunit which associates with the B-subunit. In an oxidizing environment they are joined by a disulfide bond. When Stx binds to its cellular receptor, globotriaosylceramide (GB_3_), it is internalized by the cell and the A1 and A2 subunits dissociate through reduction of the disulfide bond. The A1 subunit is free to diffuse through the cell and inactivate the 28S rRNA subunit of the ribosome via its *N*-glycosidase activity, causing protein synthesis inhibition and subsequent cell death [5].

Specifically, Stx causes removal of a catalytic adenine from the 28S rRNA, leading to the inactivation of the ribosome [6]. Stx1 and Stx2 are very similar in terms of their structure and binding to the GB_3_ receptor. However, they are trafficked through the cell via differing mechanisms that are not well understood. Additionally, Stx2 is about 1,000 times more potent than Stx1 [5].

Human *E. coli* O157:H7 infections are not well modeled in animals. For example, mice are resistant to O157:H7 infection, although they are sensitive to intraperitoneal Stx injections. To better understand how this pathogen affects humans, we have used human intestinal organoids (HIOs) and human intestinal enteroids (HIEs) to model O157 infection. HIOs are derived from pluripotent stem cells and can be differentiated into the intestinal epithelium and mesenchyme. They are grown as three-dimensional structures which recapitulate the intestinal architecture, including the lumen, villi, brush border, and crypts [7]. HIEs are derived from multipotent stem cells and have the capacity to only express the cells of the intestinal epithelium. HIEs can be grown as three-dimensional spheres, or HIEs can be processed and plated in transwell plates to grow human intestinal enteroid monolayers (HIEMs) with apical actin and basolateral nuclei. The cells can be propagated as stem cells or differentiated cells depending on the growth medium. Differentiated HIEMs possess enterocytes, goblet cells, Paneth cells, and enteroendocrine cells [7]. Once the epithelial barrier has formed, saline can be added to the apical surface to better recapitulate the intestinal environment in which the intestinal cells receive nutrients from the basolateral surface rather than the apical surface [8].

HIEMs and organoids receive different media types to support their growth. HIEMs are grown in differentiation medium while organoids are grown in organoid medium. We utilize media from StemCell Technologies, the components of which are proprietary. However, Zou et. al. [9] defines differentiation medium and organoid medium as having the same base components, and differentiation medium has Noggin, [Leu15]-Gastrin I, and A-83-01 added (Table 1). Therefore, differentiation medium will be referred to as medium with added growth factors (+GF) and organoid medium will be referred to as medium without added growth factors (no GFs added).

**Table 1:** Medium components [9].

| **Differentiation medium**<br>**(HIEMs)** | **Organoid medium (HIOs)** |
| --- | --- |
| Advanced DMEM/F12 | Advanced DMEM/F12 |
| L-Glutamine and/or Glutamax-1 | L-Glutamine |
| HEPES | HEPES |
| Pen/Strep | Pen/Strep |
| EGF | EGF |
| B27 | B27 |
| N2 | N2 |
| *Noggin | --- |
| *[Leu15]-Gastrin I | --- |
| *A-83-01 | --- |

\*Added growth factors present in differentiation medium, but not organoid medium.

Intestinal mesenchymal cells supply the epithelial cells with many different mediators to direct the growth and differentiation of stem cell populations within the intestines. Some of these factors include, but are not limited to, Wnt, BMP, and Wnt antagonists. The production of these growth factors is critical in maintaining the Wnt/b-catenin signaling pathway [11]. We aimed to determine if mesenchymal cells are able to maintain the epithelial cell monolayer when added growth factors aren’t provided in the medium, similar to how the epithelial cells and mesenchymal cells interact in organoids, where added growth factors are not provided in the medium.

Previous studies have shown that HIOs with the surrounding mesenchyme are greatly impacted by short term exposure to Stx2 compared to epithelial cells plated as HIEMs [10]. Stx2 added to the surrounding medium of HIOs or directly injected into the lumen of HIOs show loss of barrier integrity within 24-72 hours of Stx2 exposure, whereas HIEMs exposed to Stx2 for 24 hours and did not show any impact on the epithelial barrier integrity until several days post-exposure [10].

In this study, we further explored these differences and combined our HIEMs with mesenchymal cells to better recapitulate the tissue interactions between the intestinal epithelium and mesenchymal cells.

## Results

Previous studies have shown that organoids are more susceptible to Stx2 exposure through luminal or interstitial exposure. However, enteroid monolayers did not show significant damage upon Stx2 exposure [10]. This led us to hypothesize that the mesenchymal cells, which are present in organoids but not enteroids, play a role in the destruction seen during Stx2 treatment. Moreover, organoids receive a simpler medium in comparison with enteroids (Table 1), suggesting mesenchymal cells provide the epithelial cells with necessary factors for organoid survival.

### Mesenchymal cells support epithelial cell monolayer

To study this phenomenon, we combined human intestinal enteroid monolayers (HIEMs) with the mesenchymal cells from organoids to better recapitulate the tissue interactions in a transwell system. The general assay set-up can be seen in Fig. 1. We will use organoid medium, or medium without added growth factors (no GFs added), to test if the mesenchymal cells can support the epithelial cells in this transwell assay (Fig. 1). The epithelial cells include enterocytes, goblet cells, Paneth cells, and enteroendocrine cells, which are the cell types that are found in HIEMs [7].

**Figure 1:**
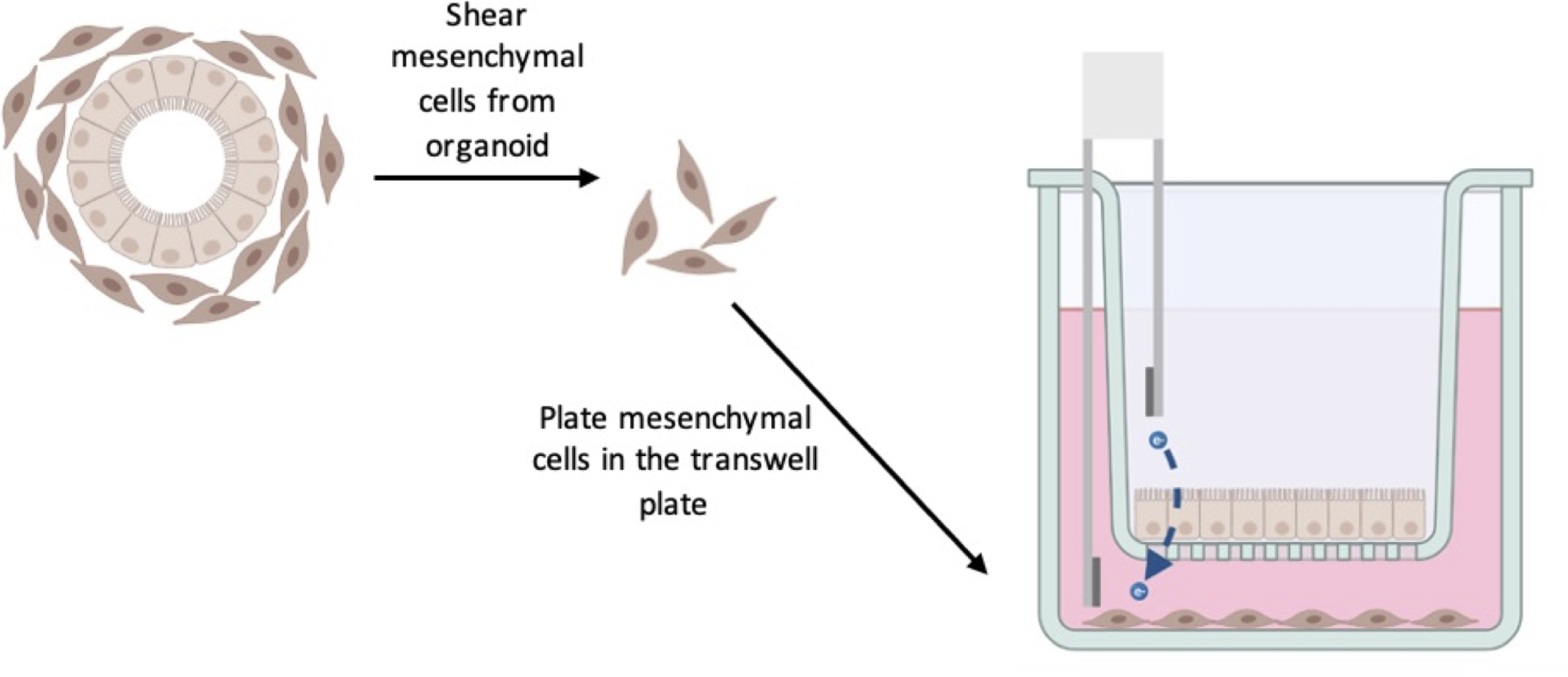
Assay set-up. Mesenchymal cells are sheared from the organoid and grown in the bottom well of a plate. Epithelial cells are grown in the transwells. Media has no growth factors added.

To test this transwell assay, epithelial cells were grown to confluency over 21 days. The apical medium was changed to saline on day 8, which was previously described in Small and Weiss [8]. On day 21, the mesenchymal cells were plated in the basolateral wells and grown with the epithelial cells for 8 days (Fig. 2A). The experiment was started by fully exchanging the spent medium for medium without added growth factors (no GFs added) or fresh medium with growth factors added (+GFs) (Fig. 2B). To prevent complete removal of putative growth factors produced by the mesenchymal cells when feeding the monolayers, a half-change of medium was performed every 3 days. To do this, 250 ml of spent medium was removed and replaced with 250 ml of fresh medium. Apical saline was fully changed every 3 days. The barrier integrity of the epithelial cell monolayer was monitored over a 12-day time course through TEER readings.

**Figure 2:**
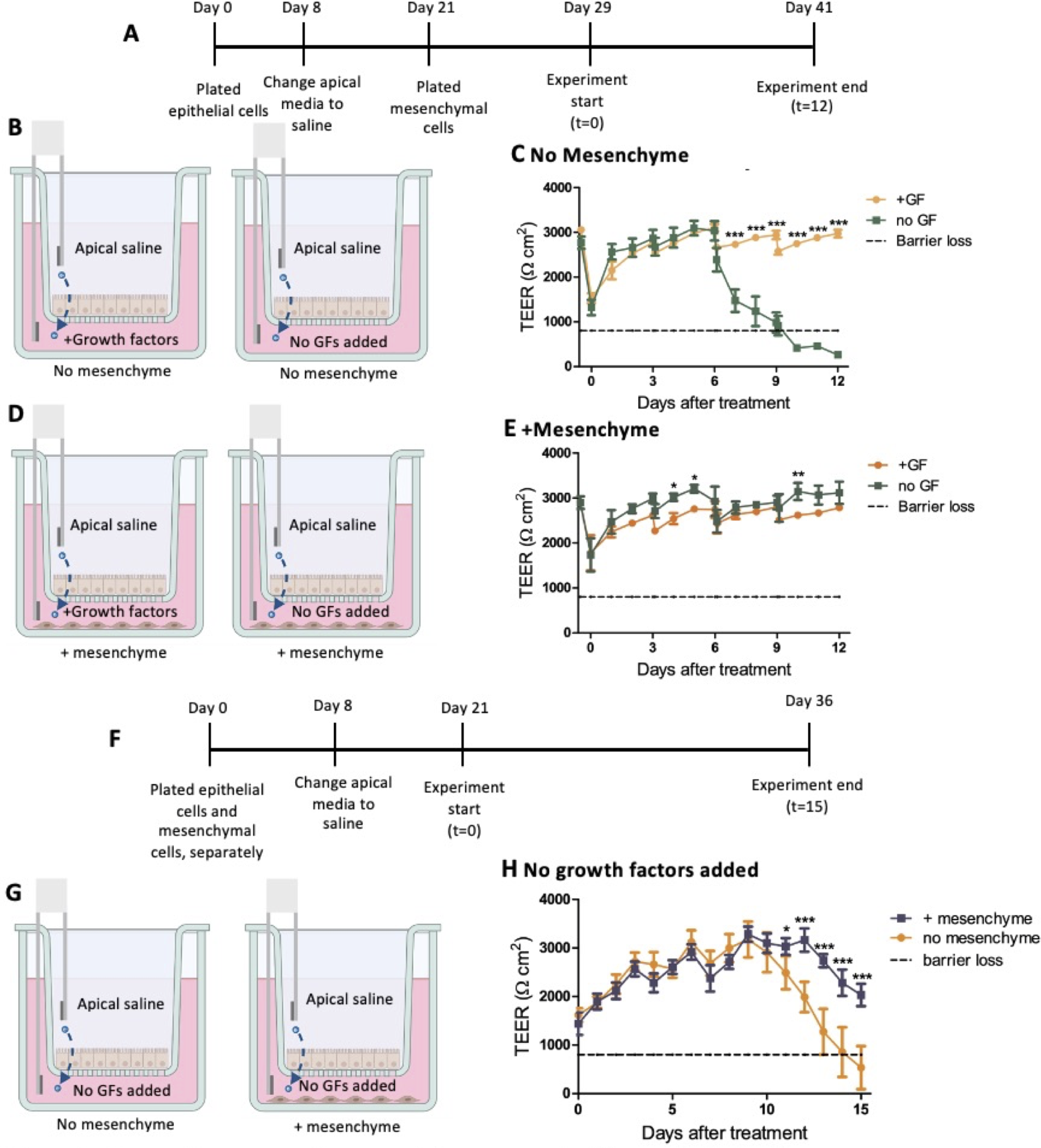
Mesenchyme supplies growth factors needed for epithelial cells. A) Timeline of experimental set-up. Mesenchymal cells were added to the basolateral wells of the transwell plates on day 21 of monolayer growth. On day 29 of monolayer growth, the media conditions were changed to either added growth factors (+GF) or no added growth factors (no GF). B) experimental conditions for no mesenchyme plate C) TEER measurements were taken over a 12-day time course; no mesenchyme, +/- GFs. D) experimental conditions for mesenchyme plate E) TEER measurements were taken over a 12-day time course; mesenchyme, +/-GFs. F) timeline of experimental set-up. Mesenchymal cells were plated at the same time and separate from the epithelial cells. The transwells were moved into the wells with the mesenchymal cells at the start of the experiment. G) Experimental conditions. H) TEER results taken over the 15-day time course. (N=3) Statistical analysis performed on GraphPad Prism, two-way ANOVA.

Without mesenchyme present in the basolateral wells, loss of TEER was noted around day 7 when added growth factors were not supplied in the medium (Fig. 2C). Alternatively, when mesenchymal cells were plated in the basolateral well (Fig. 2D), no loss of TEER was noted without added growth factors in the medium compared to wells with growth factors added (Fig. 2E). The loss of barrier integrity, shown by the dotted line, was set at 800 Ω*cm^2^, the value for a Caco2 monolayer according to Bock et. al., so lower than 800 Ω*cm^2^ would indicate loss of barrier integrity [12]. These results demonstrate that mesenchymal cells promote health of the epithelial monolayer in the absence of added growth factors. It is of note that in Fig. 2C, TEER was the same in the presence and absence of growth factors until day 7, when TEER abruptly decreased in the absence of growth factors.

To test another method of cell culture, the mesenchymal cells and the epithelial cells were grown separately for 21 days, then plated together at the start of the experiment by transferring half of the transwells to the wells with the plated mesenchymal cells (Fig. 2F and 2G). Full saline and half medium changes were performed every three days. Similar results were noted, where the mesenchymal cells supported the epithelial cells when added growth factors were not provided by the medium (Fig. 2H). Since either method of plating mesenchymal cells and epithelial cells works for this assay, both methods were used throughout the experiments described here.

### More frequent changes of media eliminate the protective effect of mesenchyme

To determine the frequency of media changes needed to maintain the epithelial barrier in this assay, we performed daily half-media changes (Fig. 3). Epithelial cells and mesenchymal cells were grown separately for 21 days (Fig. 3A). At the experiment’s start, the medium in the wells with only epithelial cells was changed to fresh medium with growth factors added or medium without added growth factors (Fig. 3B). Full apical saline and half medium changes were performed daily (Fig. 3A). TEER results showed that without added growth factors, the epithelial barrier integrity is eventually lost (Fig. 3C). Additionally at the experiment start, epithelial cells growing in the transwells were moved to wells with mesenchymal cells and medium without added growth factors. Half-medium changes and full apical saline changes were performed daily (Fig. 3A). TEER results show that barrier function sharply declined around eleven days after the start of the experiment (Fig. 3E). This demonstrates the need for longer time periods between media changes. We hypothesize that the factors being produced by mesenchymal cells accumulate in the media, reaching concentrations that are necessary to maintain epithelial barrier integrity, a result not seen with daily medium changes.

**Figure 3:**
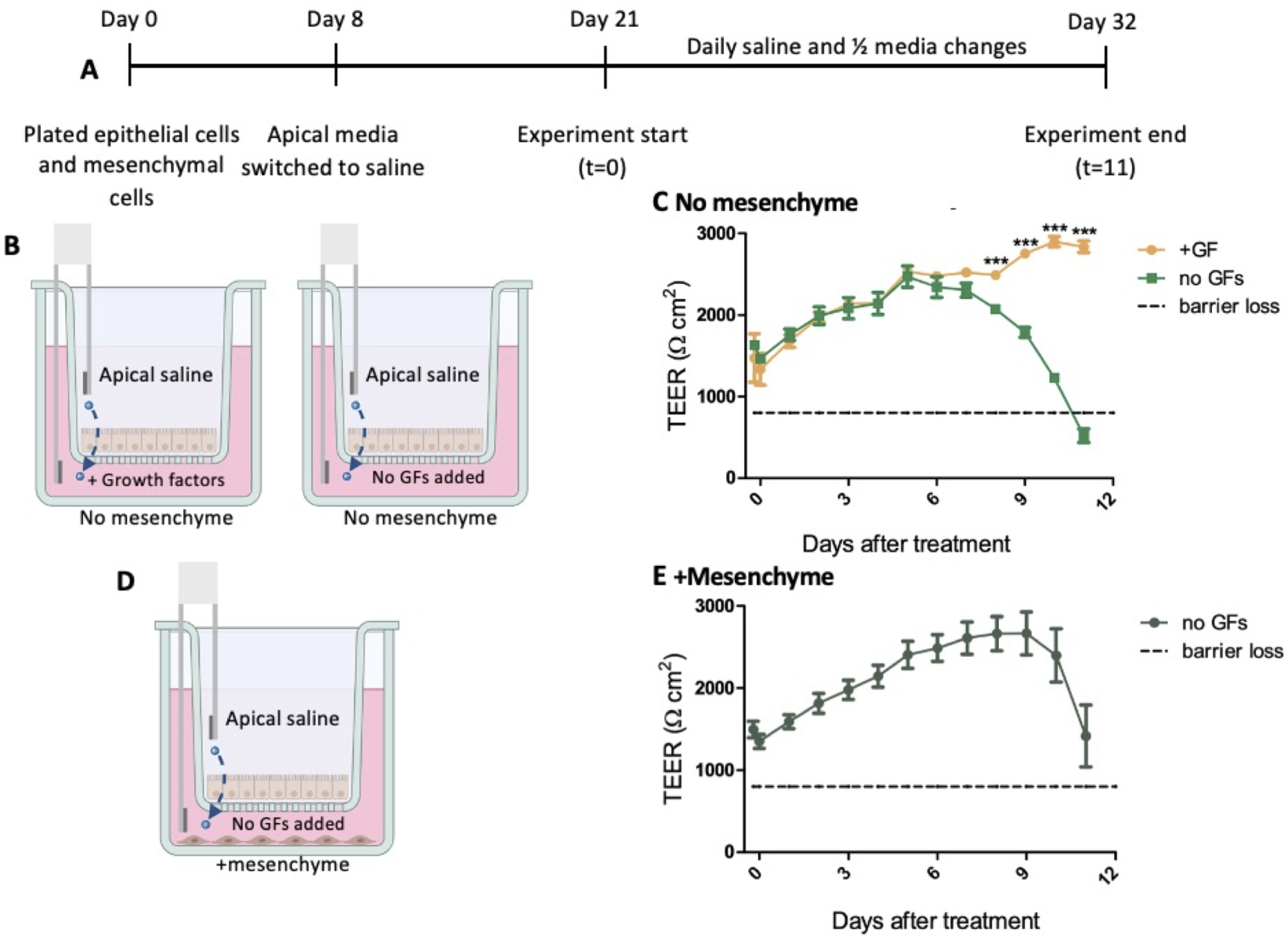
Daily medium changes lead to loss of barrier integrity. A) mesenchymal cells were plated at the same time as the epithelial cells in separate wells in transwell plates. On day 21 of monolayer growth, the apical wells with epithelial cell monolayers were transferred to the wells growing mesenchymal cells. The experimental conditions were then switched to fresh apical saline, while basolateral medium was either switched to medium with added growth factors or kept as medium without added growth factors. Apical saline was fully changed, and basolateral medium was half-changed every day. B) No mesenchyme experimental conditions. C) TEER measurements were taken over an 11-day time course. D) +mesenchyme experimental conditions. E) TEER measurements were taken over an 11-day time course. (N=3) Statistical analysis performed on GraphPad Prism, two-way ANOVA.

### Apical saline promotes barrier integrity compared to apical medium

Previous studies have shown that apical saline can be used when growing HIEMs instead of apical medium [8]. We wanted to test apical saline versus apical medium (without added growth factors) in this system. To perform this experiment, epithelial cells and mesenchymal cells were grown separately for 21 days. In the epithelial cell transwells, the apical medium was exchanged for apical saline 8 days after plating the cells (Fig. 4A). At the experiment start, the apical saline was changed to apical medium without added growth factors or fresh apical saline. The basolateral medium was changed from medium with growth factors to medium without growth factors (Fig. 4B). In contrast to the wells with apical saline, TEER readings from the wells with apical medium were inconsistent, and eventually experienced loss of barrier function (Fig. 4C). These results show that apical saline maintains the monolayer in this assay while apical medium without added growth factors does not.

**Figure 4:**
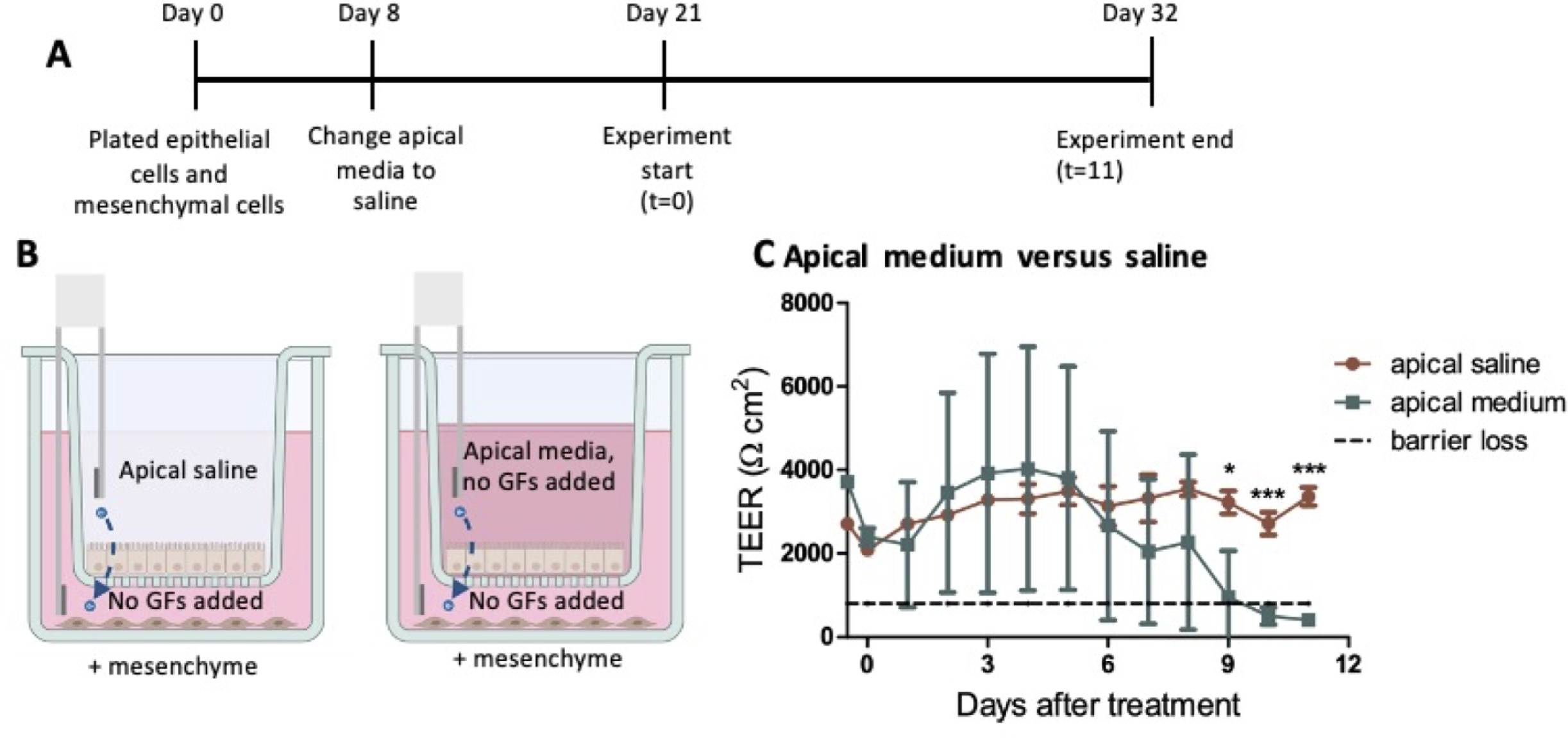
Apical saline prevents barrier loss. A&B) Mesenchymal cells were plated at the same time as the epithelial cells in separate wells on the transwell plates. On day 21 of monolayer growth, the apical wells with epithelial cell monolayers were transferred to the wells growing mesenchymal cells. The experimental conditions were then switched to apical media (no growth factors added) or fresh apical saline. Apical saline and media were changed every 3 days while basolateral media (no growth factors added) was half changed every three days (250μl of spent media was removed and replaced with 250μl of fresh media). C) TEER results were taken over an 11-day time course. (N=3) Statistical analysis performed on GraphPad Prism, unpaired t-test performed on individual timepoints.

### Mesenchymal cells are killed by medium with added growth factors

Our observations suggested that mesenchymal cells were negatively impacted by using medium with added growth factors. To test this observation, we performed an experiment in which mesenchymal cells alone were grown in medium with or without added growth factors. After 21 days of growth (Fig. 5A), mesenchymal cells were exposed to medium with added growth factors or media without added growth factors (Fig. 5B). Cells were stained with Hoechst, to stain the nuclei of living cells, and Sytox green, to stain dead cells, and observed using fluorescent microscopy at 1, 4, and 7 days (Fig. 5C&D). Sytox fluorescence shows increased cell death in wells exposed to media with added growth factors (Fig. 5E) and cell counts of live cells show a marked decrease in live cells present under medium with added growth factors condition (Fig. 5F) especially by 168 hours. These results demonstrate that medium with added growth factors does lead to mesenchymal cell death.

**Figure 5:**
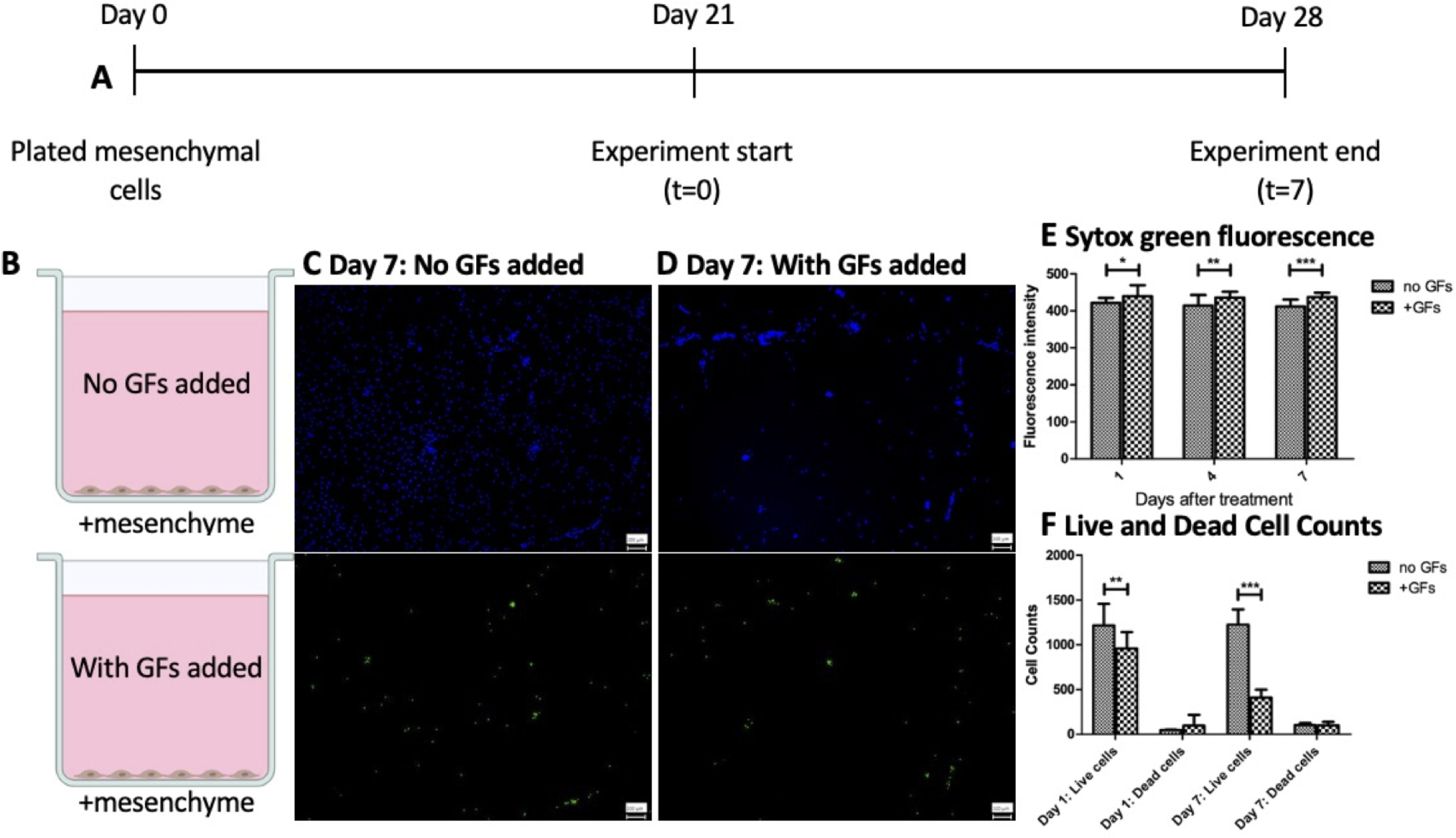
Mesenchymal cells are killed by media with added growth factors. A) mesenchymal cells were plated in 24-well plates. B) On day 21 of mesenchymal cell growth, the medium was switched to C) medium without added growth factors or D) kept as medium with added growth factors. Cells were stained with Hoechst (blue) to indicate live cells and Sytox (green) to measure dead cells. E) Fluorescence intensity was measured using ImageJ on images taken on days 1, 4, and 7. F) Cell counts of live and dead cells were performed on images taken at 1 and 7 days (N=3). Statistical analysis performed on GraphPad Prism, two-way ANOVA comparing fluorescence intensities (E) and live and dead cell counts (F).

### Apical versus basolateral addition of Stx2 is different when mesenchyme is present

Previous studies determined that a single 30 ng dose of Stx2 did not cause significant damage to HIEMs when added for 24 hours but did significantly impact HIOs [10]. We examined the impact of a high, 24-hour dose of Stx2 on the epithelial and mesenchymal cell transwell assay when added basolaterally. Epithelial cells and mesenchymal cells were grown separately for 21 days, then epithelial cell transwells were moved to mesenchymal cell wells at the start of the experiment. Mesenchymal cells were grown in medium without added growth factors (Fig. 6A). The experimental conditions were apical saline, plus a one-time dose of 30 ng Stx2 or 200 ng of Stx2, added to the apical surface, or 200 ng Stx2 added to the basolateral surface (Fig. 6B). Stx2 exposure on the apical surface was limited to 24 hours, at which time the apical saline was fully changed to fresh saline and the basolateral medium was half-changed with fresh medium without added growth factors.

**Figure 6:**
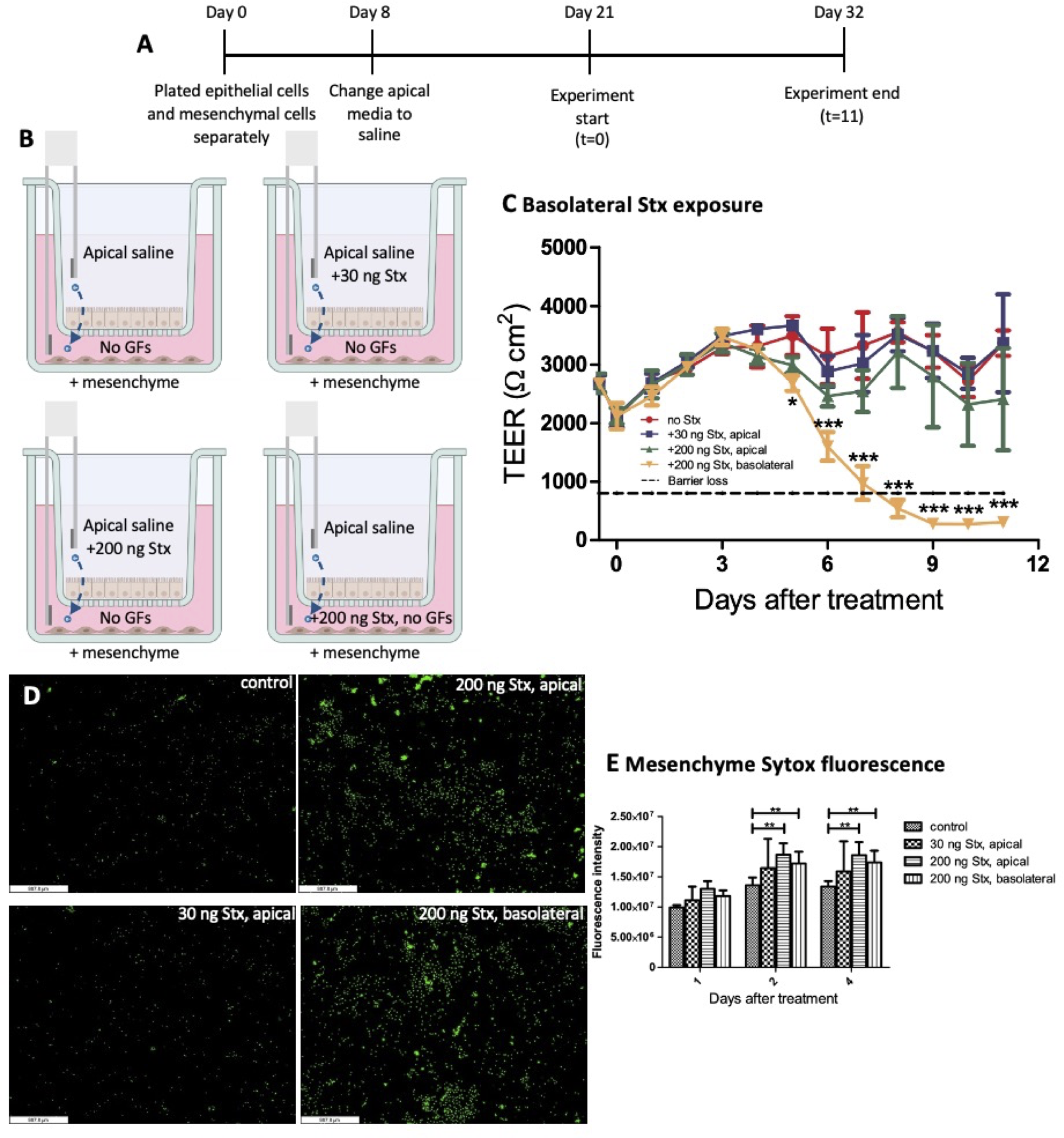
High dose of Stx induces loss of mesenchymal cells, and barrier destruction, when added basolaterally. A&B) Mesenchymal cells were plated at the same time as the epithelial cells in separate wells on the transwell plates. On day 21 of monolayer growth, the apical wells with epithelial cell monolayers were transferred to the wells growing mesenchymal cells, and fresh apical saline was added. Stx was added to appropriate wells in the apical saline or basolateral media for 24 hours. After 24 hours of exposure, apical saline was removed and replaced with fresh saline without Stx added and basolateral media was half-changed. Apical saline was changed every three days. Half of the basolateral media (no growth factors added) was changed every three days to fresh media, no Stx added. C) TEER measurements were taken over an 11-day time course. D) Mesenchymal cells were stained with Sytox green. E) Images were taken at 1-, 2-, and 4-days post-treatment and fluorescence was measured using ImageJ. (N=3) Statistical analysis performed on GraphPad Prism, two-way ANOVA.

The results indicate that a single dose of Stx2 in saline of either 30 ng or 200 ng on the apical surface did not significantly affect the barrier function. However, basolateral addition of 200 ng of Stx2 significantly disrupted the epithelial barrier integrity starting at day 5 of the time course when compared to the other treatments (Fig. 6C). Of note, when basolateral Stx2 was added, TEER was unaffected until day 6 when there was a rapid drop in TEER to below the level where a functional barrier is present.

Sytox green staining of the mesenchymal cells (Fig 6D) was performed to determine mesenchymal cell death in these conditions. Increased cell death was seen with the high levels of Stx2 treatment (200 ng) added apically or basolaterally (Fig 6E). This is in contrast to TEER readings where significant loss was only seen with basolateral treatment of Stx2 (Fig 6C). Thus basolateral addition of 200 ng of Stx2 affects the mesenchyme more than the epithelial cell layer.

### Epithelial cells alone are less susceptible to basolateral Shiga toxin exposure

To determine if the presence of mesenchymal cells drives the loss of TEER seen in Fig. 6 from basolateral Stx2 exposure or if epithelial cells alone are more susceptible to basolateral Stx2 exposure, we examined how the epithelial cells react to Stx2 without mesenchymal cells present. The epithelial cells were grown for 8 days with apical medium with growth factors added, which was exchanged for apical saline on day 8. On day 21 of cell culture, the experiment was started (Fig. 7A). No mesenchyme was provided and medium with growth factors added was provided basolaterally (Fig. 7B). The experimental conditions were apical saline, apical saline + 200 ng Stx2, basolateral medium + 30 ng Stx2, or basolateral medium + 200 ng Stx2 (Fig. 7B). In the absence of mesenchymal cells, the epithelial cells were resistant to Stx2 added apically or basolaterally (Fig. 7C).

**Figure 7:**
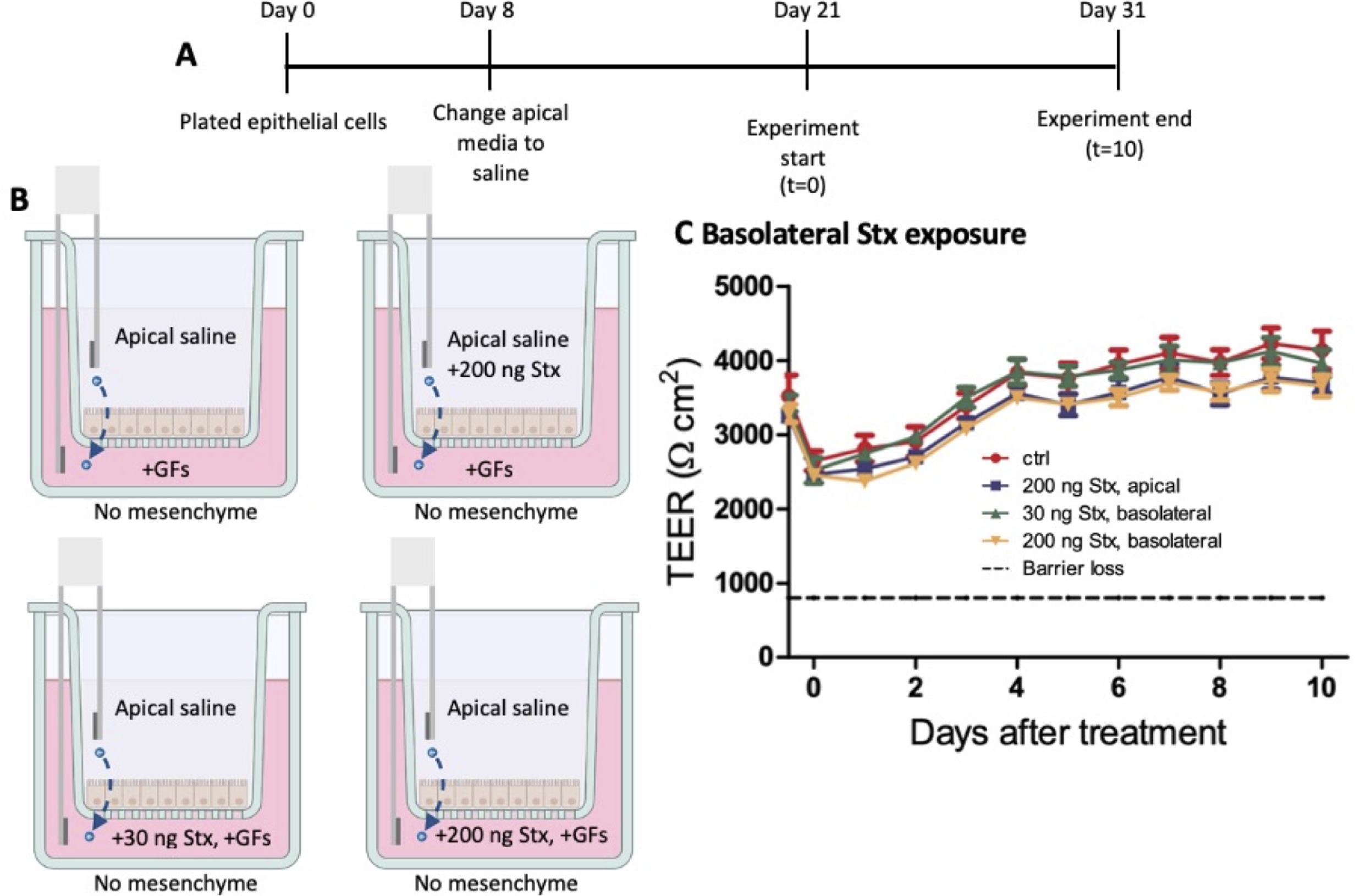
Epithelial cells without mesenchyme are not as susceptible to basolateral Stx exposure. A&B) No mesenchyme was grown in the basolateral wells for this experiment. At the experiment start, 30 ng or 200 ng of Stx was added to the apical saline or basolateral medium (with growth factors added) as appropriate. After 24 hours, apical saline was removed and replaced with fresh saline, no Stx added, and the basolateral medium was half-changed. Saline was fully changed and medium was half-changed every three days. C&D) TEER measurements were taken over a 10-day time course. (N=3) Statistical analysis performed on GraphPad Prism, two-way ANOVA, comparing controls to 200 ng Stx, basolateral.

### Repeated doses of Shiga toxin damages the epithelium

To determine the effects of Shiga toxin (Stx2) on the epithelium and mesenchyme in this system, continuous exposure of the cells to Stx2 was tested by addition of 30 ng of Stx2 to each change of apical saline at the start of the experiment and every 3 days. Epithelial cells were plated and grown for 8 days before the apical medium was exchanged for apical saline. On day 21, the mesenchymal cells were plated in the basolateral wells of half of the plates. The experiment was started 8 days later, on day 29 of epithelial cell growth (Fig. 8A). The experimental conditions were +/- 30 ng of Stx2, added to the apical saline; +/- growth factors in the basolateral medium; and +/- mesenchyme (Fig. 8B&D). The apical saline was fully changed every 3 days with 30 ng of Stx2 added to the appropriate wells. To feed the cells, half of the basolateral medium was replaced every 3 days with fresh medium, with or without growth factors added, to the appropriate wells.

**Figure 8:**
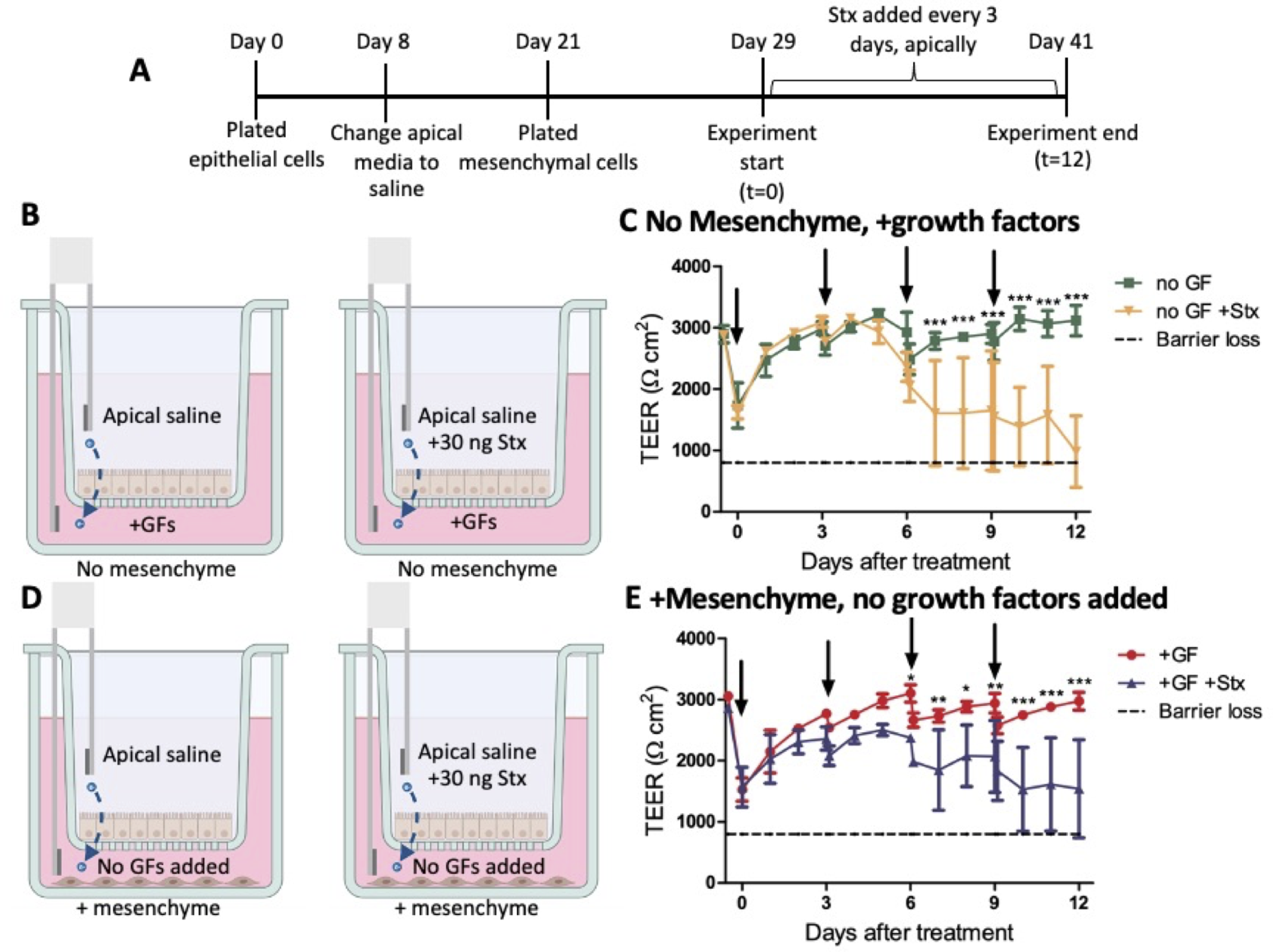
Repeated doses of Stx damages the epithelium. A) Mesenchymal cells were added to the basolateral wells of half of the transwell plates on day 21 of monolayer growth. At the start of the experiment, on day 29 of monolayer growth, the media conditions were changed to either without added growth factors (no GFs added) or added growth factors (+GFs). 30 ng of Stx2 was added to the wells as indicated at day 29 of monolayer growth. Fresh Stx2 was added to the appropriate wells at each apical saline change (every 3 days). B) experimental conditions for (C), +/- Stx2 in apical saline, with growth factors added in the medium, no mesenchyme. C) TEER measurements were taken over a 12-day time course. Arrows indicate each time Stx2 was added. D) experimental conditions for (E), +/- Stx2, no growth factors added in the medium, +mesenchyme. E) TEER measurements were taken over a 12-day time course. Arrows indicate each time Stx2 was added (N=3) Statistical analysis performed on GraphPad Prism, two-way ANOVA.

In the absence of mesenchyme with growth factors added in the medium, Stx2 caused a slight decrease in TEER (Fig. 8C), which was first seen on day seven. In the presence of mesenchymal cells and without added growth factors in the medium, continuous Stx2 exposure caused damage to the epithelial barrier, as indicated by loss of TEER (Fig. 8E). Overall, repeated doses of 30ng of Stx2 every three days lead to epithelial barrier disruption, but not complete destruction since TEER did not go below 800 Ω*cm^2^, the level that indicates loss of barrier function. These results were similar with mesenchyme or without mesenchyme in the basolateral wells.

## Discussion

These studies have shown that mesenchymal cells are needed to support epithelial cells. They act by replacing the growth factors that are otherwise present in the medium used to grow epithelial cells (Fig. 2). However, the medium with growth factors is toxic to the mesenchyme (Fig. 5). The reason for this is currently unclear.

Based on published reports by Zou, et. al. [9], differentiation medium contains noggin, [Leu15]-Gastrin I, and A-83-01. In contrast, organoid medium does not contain noggin, [Leu15]-Gastrin I, or A-83-01. These three factors comprise the major differences between organoid medium and differentiation medium, and future studies will be performed to determine which are supplied by the mesenchyme. It is important to note that our lab utilizes media from Stemcell Technologies, the formulation of which is proprietary, and therefore it is not guaranteed that these are the sole factors involved.

Apical medium has been shown to negatively affect the TEER levels in epithelial cells alone [8], or with mesenchymal cells present (Fig. 4). This is perhaps due to the addition of medium buffered at pH 7.6 in what should be an acidic compartment. We therefore used apical saline in these studies to promote monolayer maintenance.

These Stx2 studies and our previous studies [10] have demonstrated that without mesenchymal cells, a single dose of Stx2, apically, causes very little loss of TEER and does not greatly impact the epithelial barrier for many days post-exposure. Certain cell populations may be more susceptible Stx than others, which could explain the delayed disruption of the barrier integrity once the death of those cells affected the epithelium at large. Further studies are needed to determine which cell types, if any, are killed by Stx exposure. Basolateral Stx exposure does not greatly impact the epithelium alone. Repeated doses of Stx2, apically, also does not impact the epithelium for many days post-exposure but is ultimately more damaging to the epithelium than a single, short-term exposure.

While apical exposure mimics in vivo conditions where bacteria in the lumen of the intestines are producing Stx2, basolateral exposure can help to model instances where leaky intestinal epithelium allows for increased Stx exposure interstitially. Additionally, Stx2 is capable of transcytosing across the epithelium without disrupting the epithelial barrier, also allowing for interstitial exposure.

Overall, the results of these studies have cemented the need for complex cell culture models when studying host-pathogen interactions. Common animal models such as mice are resistant to *E. coli* O157:H7 infections and intestinal delivery of Stx2, while humans appear to be sensitive to both. It has been proposed that in humans, STEC-mediated intestinal damage destroys the intestinal barrier and allows basolateral access to Stx2. In mice, there is no epithelial damage, therefore they are resistant to epithelial delivery of Stx2, while remaining sensitive to Stx2 injection. However, our studies show that like mice, the human epithelial layer is quite resistant to Stx2, and it is the sensitivity of the mesenchymal cells that in turn kills the epithelial cells. We have shown in this study, and directly in a previous study [10], that Stx2 is transported through the intact epithelium without causing damage to the resistant epithelial layer. While these studies have focused on the impact of Stx2 on the intestinal epithelium and mesenchyme, this model may also be useful for future studies into the mechanisms of Stx1 transcytosis and intestinal tissue response. Understanding tissue interactions during infections is critical in determining the effects of pathogens on human tissues.

## Methods

### Maintenance of HIOs

Human intestinal organoids (H1 HIOs) were obtained from the Pluripotent Stem Cell Facility and the Organoid Core at Cincinnati Children’s Hospital and Medical Center. Organoids were grown as previously described [7]. HIOs were grown in gut medium, which is composed of DMEM/F12 (gibco, 12634-010), 1x B-27 (gibco, 12587-001), 1x N-2(gibco, 17502-001), 15 mM HEPES (Quality Biological, 118-089-721), 1x L-glutamine (VWR life science, 02-0131-0100), 50 ng/ml rhEGF (R&D, 236-EG), and 1% Penicillin Streptomycin (gibco, 15140-122)

### Growth of HIEs

Human intestinal enteroids were originally derived from the implantation of H1 HIOs in mouse kidneys and were maintained in culture suspended in Matrigel (corning, 354234) and grown in OGMH, a 1:1 mixture of organoid supplement (Stemcell technologies, 100-0191) and Intesicult™ OGM Human Basal Medium (Stemcell technologies, 100-0190).

Medium was changed every 2-3 days and enteroids were passaged after 5 days of growth to expand the culture. To passage, enteroids were passed through a 1cc U-100 insulin syringe (Becton Dickinson, 329424) 3 times to fragment the spheroids, centrifuged at 3800xg for 1 minute, then suspended in Matrigel and plated in 4-well plates. Plates were incubated upside down until the Matrigel solidified (about 15 minutes) then 500 ml of OGMH was added.

### Growth of HIEMs

Human intestinal enteroids (HIEs) were from the H1 Line (WA01) NIH Registration #0043 human embryonic stem cell line, normal 46 XY karyotype, as previously described [7]. Enteroids were processed and grown in transwells (Costar, 3470), coated with collagen (Sigma, C5533-5MG), using a modified protocol from Zou, et al [9]. Monolayers are grown in expansion medium, OGMH with 10 mM/L Y-27632 (Stemcell technologies, 72305) added. After 24 hours, the medium was changed to differentiation medium, a 1:1 mix of Intesicult™ OGM Human Basal Medium and gut medium, 200 ml apical medium and 500 ml basolateral medium. Medium changes occurred every 2-3 days. The apical medium was switched to saline after 8 days of cell growth. Medium and saline changes occurred every 2-3 days. Monolayers were used for experiments on day 21 of cell growth. TEER measurements were determined through ERS-2 volt-ohm meter (Millicell).

### Maintenance of Mesenchyme

H1 HIOs were received and grown for about 2 weeks, then mesenchymal cells were sheared from the HIOs and maintained suspended in Matrigel with 500 ml of gut medium. Medium changes occurred every 3-5 days.

### Stx2

Stx2 holotoxin (bei resources, NR-4478) was added to the apical or basolateral wells as indicated (30 ng or 200 ng).

